# A negative feedback model to explain regulation of SARS-CoV-2 replication and transcription

**DOI:** 10.1101/2020.08.23.263327

**Authors:** Shan Gao, Zhi Cheng, Xiufeng Jin, Fang Wang, Yibo Xuan, Hao Zhou, Chang Liu, Jishou Ruan, Guangyou Duan, Xin Li

## Abstract

**Background:** Coronavirus disease 2019 (COVID-19) is caused by severe acute respiratory syndrome coronavirus 2 (SARS-CoV-2). Although a preliminary understanding of the replication and transcription mechanisms of SARS-CoV-2 has recently emerged, their regulation remains unclear.

**Results:** Based on reanalysis of public data, we propose a negative feedback model to explain the regulation of replication and transcription in—but not limited to—SARS-CoV-2. The key step leading to new discoveries was the identification of the cleavage sites of nsp15—an RNA uridylate-specific endoribonuclease, encoded by CoVs. According to this model, nsp15 regulates the synthesis of subgenomic RNAs (sgRNAs) and genomic RNAs (gRNAs) by cleaving transcription regulatory sequences in the body. The expression level of nsp15 determines the relative proportions of sgRNAs and gRNAs, which in turn change the expression level of nps15 to reach equilibrium between the replication and transcription of CoVs.

**Conclusions:** The replication and transcription of CoVs are regulated by a negative feedback mechanism that influences the persistence of CoVs in hosts. Our findings enrich fundamental knowledge in the field of gene expression and its regulation, and provide new clues for future studies. One important clue is that nsp15 may be an important and ideal target for the development of drugs (e.g. uridine derivatives) against CoVs.

## Introduction

Coronavirus disease 2019 (COVID-19) is caused by severe acute respiratory syndrome coronavirus 2 (SARS-CoV-2). As enveloped viruses composed of a positive-sense, single-stranded RNAs, CoVs have the largest genomes (26–32 kb) among all RNA virus families. SARS-CoV-2 has a genome of ∼30 kb [1]. In addition to ORF1a and 1b (**Figure 1A**), the SARS-CoV-2 genome has sequences encoding four conserved structural proteins [spike protein (S), envelope protein (E), membrane protein (M), and nucleocapsid protein (N)] and six accessory proteins (ORF3a, 6, 7a, 7b, 8, and 10) that are yet to be experimentally verified.

**Figure 1.**
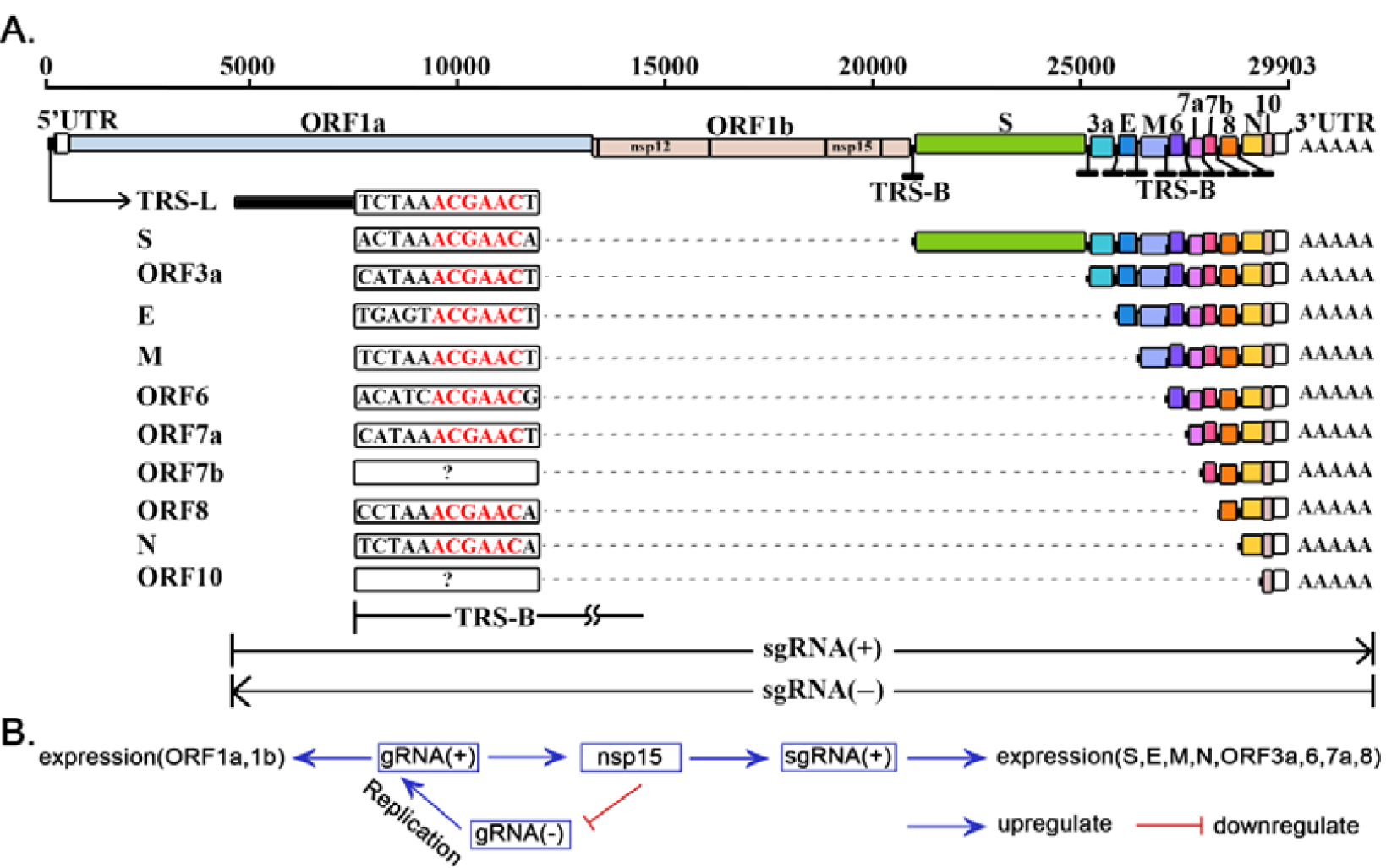
Replication and transcription of SARS-CoV-2. The elements used to represent the SARS-CoV-2 genome (GenBank: MN908947.3) were originally used in the previous study [2]. **A**. TRS-L: transcription regulatory sequence in the body; TRS-B: transcription regulatory sequence in the leader. **B**. gRNA(+):genomic RNAs; gRNAs(-): antisense genomic RNAs.

The genomic RNA, as a positive-sense RNA [gRNA(+)], is used as a template for the translation of ORF1a and ORF1b, and the replication and transcription of the SARS-CoV-2 genome. Recently, the translation of 10 other proteins (S, E, M, N, ORF3a, 6, 7a, 7b, 8 and 10) of SARS-CoV-2 has been reported in a study [2] that directly validates the prevailing “leader-to-body fusion” model using Nanopore RNA-seq—a direct RNA sequencing method [3]. In the model validated in that study (**Figure 1A**), replication and transcription require gRNA(+) as the template for the synthesis of antisense genomic RNAs [gRNAs(-)] and antisense subgenomic RNAs [sgRNAs(-)] by RNA-dependent RNA polymerase (RdRP). When RdRP pauses, as it crosses a body transcription regulatory sequence (TRS-B) and switches the template to the leader TRS (TRS-L), sgRNAs(-) are formed. Otherwise, RdRP reads gRNAs(+) continuously, without interruption, resulting in gRNAs(-). Thereafter, gRNAs(-) and sgRNAs(-) are used as templates to synthesize gRNAs(+) and sgRNAs(+), respectively; sgRNAs(+) are used as templates for the translation of the 10 proteins mentioned above.

TRS-L usually forms the first 60–70 nts of the 5’ UTR in a CoV genome. TRS-B with varied length is located immediately upstream of ORFs except ORF1a and 1b (**Figure 1A**). During antisense-strand synthesis, the discontinuous transcription (referred to as polymerase jumping or template switching) by RdRP is described by the “leader-to-body fusion” model that is conserved in the entire nidovirus order, including arteriviruses, toroviruses, roniviruses and animal coronaviruses. Although investigation was conducted to determine the underlying mechanisms in previous studies [4], the molecular basis remain unknown. In the present study, we aimed to determine the molecular basis of the “leader-body fusion” model and construct a model to explain the regulation of SARS-CoV-2 replication and transcription.

## Results

### Reanalysis of leader-body junction regions

Using public Nanopore RNA-seq data (**Materials and Methods**), 575,106 out of 879,679 reads from a SARS-CoV-2-infected Vero cell sample were aligned to the SARS-CoV-2 reference genome. Among all aligned reads, 575,106 sense reads represented gRNAs(+) or sgRNAs(+), while 30 antisense reads represented gRNAs(-) or sgRNAs(-). This exceedingly high ratio (575,106 vs. 30) between sense and antisense reads may be the result of significant differences between the gRNAs(+)/sgRNAs(+) and gRNAs(-)/sgRNAs(-) degradation efficiencies. This phenomenon, however, was not reported in the previous study [2]. Another explanation for the high ratio is that gRNAs(+)/sgRNAs(+) are protected by binding to the N proteins. Another super high ratio (198,198,542 vs. 11,820,438) between contiguous and junction-spanning reads was reported in that previous study [2]. This suggested that there were significant differences between gRNAs(+)/gRNAs(-) and sgRNAs(+)/sgRNAs(-).

By reanalysis of junction-spanning reads, we found that TRS-Bs share ∼12 nt junction regions with TRS-Ls in SARS-CoV-2 (**Figure 1A**). Most junction regions in eight protein-coding genes (*S, E, M, N, ORF3a, 6, 7a and 8*) exhibit high similarity to those of TRS-Ls and are defined as canonical junction regions [2]. Accordingly, the sgRNAs(+) and sgRNAs(-) containing these canonical junction regions are defined as canonical sgRNAs(+) and sgRNAs(-), respectively. The junction regions of *ORF7b* exhibit high diversity in their sequences. Whether junction regions exist in *ORF10* remains unclear [2]. Our analysis of canonical junction regions determined their core sequence (CS) of SARS-CoV-2 to be ACGAAC (**Figure 1A**).

### Further investigation of “leader-to-body fusion” model

Further analysis of betacoronavirus subgroup B (**Materials and Methods**) showed that ACGAAC is a highly conserved sequence among their genomes, particularly in their putative TRS-Bs. This suggests that canonical junction regions contain a specific motif for enzyme reorganization. Since all CoVs and even viruses in the entire nidovirus order adhere to the “leader-to-body fusion” model [2], this or these enzymes should be encoded by the *ORF1a* or *1b* gene, given their likelihood to be translated. After analysing 16 non-structural proteins (nsp1–16) encoded by the *ORF1a* or *1b* genes, we determined that nsp15 (a nidoviral RNA uridylate-specific endoribonuclease, NendoU [5]) is most likely to function in these junction regions, given that a homolog of nsp15 has cleavage sites containing “GU” [6]. Thus, the cleavage site of nsp15 was identified to follow the motif “GTTCGT|N” (the vertical line indicates the breakpoint and N indicates any nucleotide base), read in the antisense strands of CoV genomes. Furthermore, we found that almost all the genomic sites containing the motif “GTTCGT” have polyT (not less than three T) at the tail, which ensures the presence of at least one U for nsp5 cleavage.

Upon searching for “GTTCGT” in the genomes of betacoronavirus subgroup B, the occurrence of “GTTCGT” on the antisense strand was found to be more than 1.6 times that on the sense strand. In particular, “GTTCGT” occurred 3 and 9 times (**Table 1**) on the sense and antisense strands of the SARS-CoV-2 genome, respectively. These findings suggest that the basic function of nsp15 involves the degradation of gRNAs and gsRNAs and that the high ratio between sense and antisense reads (**see above**) results from substantially more cleavage of gRNAs(-)/gsRNAs(-) than that of gRNAs(+)/gsRNAs(+). Among the three sites containing “GTTCGT” on the sense strand of the SARS-CoV-2 genome (referred to as internal cleavage sites—ICSs), one is located in the coding sequence (CDS) of RdRP (nsp12), while the other two are located in the *ORF8* gene (**see below**).

**Table 1.** The genomic sites of motif GTTCGT. The motifs (in the first column) were mapped to the SARS-CoV-2 genome (GenBank: MN908947.3) using positions (in the second column) of the first nucleotides. ACGAAC indicated the motif GTTCGT, read on the antisense strand of the SARS-CoV-2 genome. The third column shows the regions influenced by the motifs (in the first column). * this site is located in the CDS of RdRP (nsp12). TRS-L: transcription regulatory sequence in the body; TRS-B: transcription regulatory sequence in the leader; ICS: internal cleavage site; RdRP: RNA-dependent RNA polymerase.

| Motif | Position | Region(Start-end) | Type |
| --- | --- | --- | --- |
| GTTCGT | 16014 | nsp12(13483-16236)* | ICS |
|  | 28198 | ORF8(27894-28259) | ICS |
|  | 28233 | ORF8(27894-28259) | ICS |
| ACGAAC | 70 | 5' UTR(1-265) | TRS-L |
|  | 21556 | S(21563-25384) | TRS-B |
|  | 25385 | ORF3a(25393-26220) | TRS-B |
|  | 26237 | E(26245-26472) | TRS-B |
|  | 26473 | M(26523-27191) | TRS-B |
|  | 27041 | ORF6(27202-27387) | TRS-B |
|  | 27388 | ORF7a(27394-27759) | TRS-B |
|  | 27888 | ORF8(27894-28259) | TRS-B |
|  | 28260 | N(28274-29533) | TRS-B |

These two ICSs are also located in *ORF8* of most SARS-CoV-2, SARS2-like CoV and SARS-like CoV genomes; however, they are absent in the genomes of SARS-CoVs obtained from humans (GenBank: AY274119 and AY278489) and SARS-like CoV genomes from civets (GenBank: AY304486, AY515512 and AY572034). One of the two ICSs is present in the genome of the SARS-like CoV strain WIV1 from bats (GenBank: KF367457), which is the ancestor of SARS-CoV. Deletions of *ORF8* were reported to be associated with attenuation of SARS-CoV (GenBank: AY274119) [8] and SARS-CoV-2 [10]. The *ORF8* gene of SARS-CoV is considered to have played a significant role in adaptation to human hosts following interspecies transmission [7] via the modification of viral replication [8]. The loss of two nsp15 ICSs in ORF8 of SARS-CoV is the key clue revealing the functions of ORF8 and the pandemic of SARS-CoV.

Based on the above analyses, we modified the “leader-body fusion” model and proposed the molecular basis of it. In our model, nsp15 cleaves synthesized gRNAs(-) and sgRNAs(-). The cleavage occurs at TRS-Bs(-) synthesized using TRS-Bs as templates. Next, the free 3’ ends (∼6 nts) of TRS-Bs(-) hybridize the junction regions of TRS-Ls for template switching. Alternatively, the cleaved TRS-Ls(-) are used as templates to synthesize TRS-Ls that then hybridize the junction regions of the cleaved sgRNAs(-) without TRS-Ls(-) for template switching and gRNAs(+)/sgRNAs(+) synthesis. These findings suggests that RdRP (nsp12) is so active that it must have a very high enzyme activity.

Some non-canonical sgRNAs(+) and sgRNAs(-) are synthesized as a result of occasional hybridization between the free 3’ ends of TRS-Bs(-) and highly similar sequences of TRS-L junction regions. The sgRNAs(-) without TRS-Ls are also synthesized due to missing hybrids. In the previous study [2], sgRNAs(-) without TRS-Ls, non-canonical sgRNAs(+) and sgRNAs(-) were reported, supporting our model. Furthermore, cleavage also occurs at ICSs on the sense strands, after which the free 3’ ends of the cleaved 5’ ends hybridize the antisense strands to synthesize recombinant sgRNAs(+) or even gRNAs(+). This may also contributes to the multiple recombination events in betacoronavirus genomes.

### Proposal of a negative feedback model

Based on the above analyses, we proposed a negative feedback model (**Figure 1B**) to explain the regulation of replication and transcription in—but not limited to—CoVs. In this model (**Figure 1**), nsp15 regulates the synthesis of subgenomic RNAs (sgRNAs) or genomic RNAs (gRNAs) by the cleavage of TRS-Bs(-). The expression level of nsp15 determines the relative proportions of sgRNAs and gRNAs. An increase in nsp15 expression results in less gRNAs(-) and more gsRNAs(-), after which fewer gRNAs(+) and more gsRNAs(+) are synthesized, respectively. A decrease in gRNAs(+) results in a decrease of nsp15 expression, given that nsp15 is synthesized using gRNA(+) as the template. Furthermore, the nsp15 ICS in the CDS of SARS-CoV-2 nsp12 (**Table 1**) enhances the negative feedback. Via this negative feedback mechanism, CoVs reach equilibrium between the replication and transcription (**Figure 1B**); thus, this mechanism is important for the persistence of CoVs in hosts.

Our negative feedback model is based on the determination of the molecular basis of the “leader-body fusion” model. This molecular basis is associated to the cleavage function of nsp15, which is different from other models proposed in the previous studies [4], mainly due to the integration of information from many aspects, particularly considering: (1) The direct RNA sequencing data [2]; (2) The nsp15 structure in complex with GpU (pdb code: 6×1B); (3) the polyT at the tail of “GTTCGT”; and (4) the nsp15 ICSs in *ORF8* (**see above**). These discoveries confirmed the identification of the cleavage sites of nsp15. However, they were not used to construct the models in the previous studies [4].

### The necessity of negative feedback regulation

In our previous study [7], we proposed that first hairpins (immediately upstream of ORF1a) have an important role in the functions (e.g. regulation of translational initiation) of ribosome binding sites (RBSs) in 5’ UTRs of the SARS-CoV-2 genomes. SARS-CoV and SARS-CoV-2 have an identical first hairpin, which may enhance the translation of downstream genes. To indirectly prove that negative feedback is a basic mechanism, we designed preliminary experiments to show that first hairpins from SARS-CoV-2 enhance the translation of the downstream enhanced green fluorescent proteins (EGFPs); however, over-expression of EGFP without negative feedback regulation will cause cell death.

In total, three types of plasmids containing EGFP reporter genes—named pEGFP-C1, pSARS, and pCoV-ba (betacoronavirus subgroup A)—were used in the experiments (**Figure 2A**). The plasmid pEGFP-C1 was used as a control, given that it contains 17-nt sequences, encoding the first hairpin from Cytomegalovirus (CMV). Two types of plasmids proceeded by 30-and 29-nt inserts were used to evaluate their influence on translation (**Materials and Methods**). These two inserts encoded the two first hairpins from SARS-CoV-2 and the subgroup A of betacoronaviruses, respectively. Comparing the fluorescent brightness of cells transfected with three types of plasmids, the expression of EGFP in pSARS was markedly higher than that in pEGFP-C1 and pCoV-ba (**Supplementary 1**). Moreover, pSARS caused cell death at 48 hours after plasmid transfection. We then performed 3-(4,5-dimethyl-2-thiazolyl)-2,5-diphenyl-2-H-tetrazolium bromide (MTT) and lactate dehydrogenase (LDH) experiments to further evaluate the influence of plasmid transfection (**Materials and Methods**). Both MTT and LDH experiments consistently suggested that pSARS caused significantly more HEK293T and Hela cell death at 48 hours after plasmid transfection, than pEGFP-C1 and pCoV-ba. However, this phenomenon appeared in HEK293 cells at 56 hours after plasmid transfection (**Figure 2B**). Given that the only difference among the three types of plasmids is their 17-, 30-, and 29-nt sequences encoding different hairpins, we concluded that these hairpins determined the translation efficiency of the downstream EGFPs. The hairpin in pSARS resulted in the over-expression of EGFP, which caused more cell death. To determine whether the factor acts at the translation level and to rule out other possible factors that may exert influence at the replication or transcription level, we performed the following experiments: (1) using HEK293 cells to rule out the possible influence by the differences of plasmid copy numbers, since all three types of plasmids containing the SV40 origins can be replicated to a copy number of between 400∼1000 plasmids per cell within HEK293T; and (2) using qPCR to rule out the possible influence by differential transcription (**Supplementary 1**).

**Figure 2.**
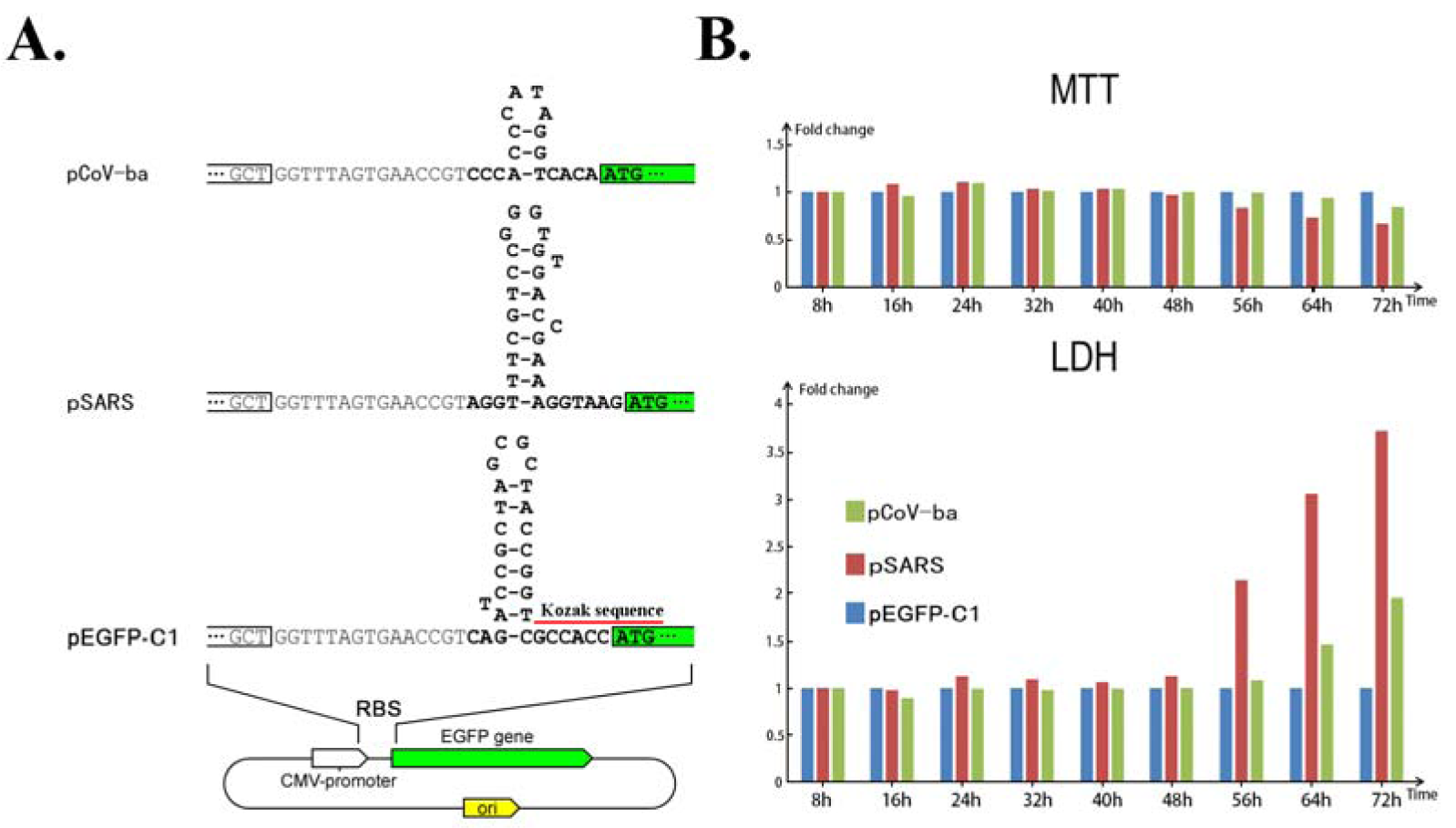
Regulation of EGFP expression by three RBSs. **A**. The plasmid pEGFP-C1 was used as control, since it contains 17-nt sequences, encoding the first hairpin from Cytomegalovirus (CMV). Two other types of plasmids pSARS and pCoV-ba were proceeded by replacement of first hairpins from SARS-CoV-2 and betacoronavirus subgroup A, respectively (**Materials and Methods**). **B**. HEK293 cells were transfected with three types of plasmids. At each time after transfection, cell viability and cell death were meausured using MTT and LDH, respectively.

## Conclusion and Discussion

In the present study, we determined the molecular basis of the “leader-body fusion” model and constructed a negative feedback model to explain the regulation of SARS-CoV-2 replication and transcription. The key step that led us to new discoveries was our identification of the cleavage sites of nsp15, an RNA uridylate-specific endoribonuclease (NendoU). Our model does not rule out other RNAs or proteins involved in these two models. However, current studies involving NendoU remain contradictory in terms of their findings regarding fundamental questions [5]: (1) NendoU is conserved among coronaviruses, arteriviruses and toroviruses, is it present in nonvertebrate-infecting representatives of the nidoviruses order? (2) is nsp15 indispensable in viral replication and living? (3) is nsp15 responsible for protein interference with the innate immune response? and (4) does nsp15 degrade viral RNA to hide it from host defences? Nsp15 cleavage sites in relevant genes provide new clues to answer these questions.

The Kozak consensus sequence GCCACCAUGG (**Figure 1B**) is not necessary for protein translation in eukaryotes. The hairpins immediately upstream of ORF1a (first hairpins) play an important role in the functions of CoV RBSs. Hairpin with proper structures may enhance the translation of their downstream proteins. These findings can be used to design vaccine, drug and expression vectors. Based on our preliminary experiments, we concluded that the hairpin in pSARS resulted in the over-expression of EGFP, which caused cell death. These facts supported that the negative feedback is necessary to prevent the over-expression of viral genes.

Our findings enrich fundamental knowledge in the field of gene expression and its regulation, and provide new clues for future studies. The template switching ability and the high ratio between contiguous and junction-spanning reads indicated that RdRP (nsp12) has a very high enzyme activity. Relatively few junction-spanning reads indicated that nsp15 does not have a high enzyme activity. This suggested that nsp15 may be an important and ideal target for the development of drugs against CoVs. The most recent nsp15 structure in complex with GpU (PDB: 6×1B) shows that the uridine in GpU binds to the active site of nsp15. Thus, uridine derivatives, such as Trifluridine, Tipiracil, Tegafur, Carmofur, Furtulon, etc, are potential inhibitors of this enzyme.

## Materials and Methods

1,265 genome sequences of betacoronaviruses were downloaded from the NCBI GenBank database in our previous study [8]. Nanopore RNA-seq data was downloaded from the website (https://osf.io/8f6n9/files/) for the reanalysis of leader-body junction regions. The results were confirmed using Illumina RNA-seq data from the NCBI SRA database under the accession number SRP251618. Data cleaning and quality control were performed using Fastq_clean [9]. Statistics and plotting were conducted using the software R v2.15.3 with the package ggplot2 [10]. The 5’ and 3’ ends of gRNAs(+) and sgRNAs(+) were observed and double-checked using the software Tablet v1.15.09.01 [11].

In the present study, three types of plasmids pEGFP-C1 (maintained in the lab of Hao Zhou), pSARS and pCoV-ba, and three types (Hela, HEK293 and HEK293T) of cell maintained in our lab were used for transfection. To construct pSARS, pEGFP-C1 was PCR amplified, using primers fVR and rRBS2 (**Supplementary 1**). Then the linear PCR product was self-ligated into a plasmid by homologous recombination technology using ClonExpress II One Step Cloning Kit (Vazyme Biotech, China). Following the same procedure, primers fVR and rRBS3 (**Supplementary 1**) were used to construct pCoV-ba. The cells were cultured in Dulbecco’s Modified Eagle Medium (DMEM) media supplemented with 10% fetal bovine serum. For each experiment, 100,000 cells were seeded into one well of a 6-well plate for plasmid transfection. After 12 hours (0 hour in **figure 2B**), transfection of 1 μg of plasmid into one well was performed using 3 μL PolyJet (SignaGen Laboratories, USA), following the manufacturer’s instructions. The medium was changed at 12 hours after plasmid transfection. MTT (200 μl, 5 mg/ml), and 1.8 ml medium was added to each well and cultured at 37 °C for 4 h. Then, the cells in each well were removed of medium, washed with PBS, then mixed with 1000 μl DMSO to dissolve the formazan product. Finally, formazan absorbance was measured by a microplate reader with a wavelength of 492 nm (Thermo Labsystems Helsinki, Finland). LDH experiments were performed using LDH cytotoxicity assay detection kit (Beyotime, China), following the manufacturer’s instructions.

## Supplementary information

Additional file 1: Figure S1. Comparison of fluorescent brightness. Table S1. Primers for plasmid construction.

## Abbreviations

COVID-19: Coronavirus disease 19
CoVs: coronavirus
SARS-CoV: Severe Acute Respiratory Syndrome Coronavirus
SARS-CoV-2: SARS Coronavirus 2
SARS-like CoV: SARS-CoV like Coronavirus
SARS-2 like CoV: SARS-CoV-2 like Coronavirus
gRNA: genomic RNA
sgRNA: subgenomic RNA
RBS: ribosome binding site
ORF: open reading frame
CDS: coding sequence
TRS-L: transcription regulatory sequence in the leader
TRS-B: transcription regulatory sequence in the body
ICS: internal cleavage site
RdRP: RNA-dependent RNA polymerase
nsp: non-structural protein
NendoU: nidoviral RNA uridylate-specific endoribonuclease
CMV: Cytomegalovirus
MTT: 3-(4,5-dimethyl-2-thiazolyl)-2,5-diphenyl-2-H-tetrazolium bromide
LDH: lactate dehydrogenase

## Declarations

### Ethics approval and consent to participate

Not applicable.

### Consent to publish

Not applicable.

### Availability of data and materials

All data used in the present study was download from the public data sources.

### Authors’ contributions

Shan Gao conceived the project. Xin Li and Guangyou Duan supervised this study. Guangyou Duan and Xiufeng Jin conducted programming. Zhi Cheng, Hao Zhou and Fang Wang conduced the experiments. Shan Gao and Yibo Xuan downloaded, managed and processed the data. Shan Gao drafted the main manuscript text. Shan Gao, Chang Liu and Jishou Ruan revised the manuscript.

## Acknowledgments

We are grateful for the help from the following faculty members of College of Life Sciences at Nankai University: Xuetao Cao, Deling Kong, Quan Chen, Wenjun Bu, Ting Ma, Tao Zhang, Dawei Huang, Mingqiang Qiao, Yanqiang Liu, Qiang Zhao, Bingjun He and Zhen Ye. This manuscript was online as a preprint on Aug 24nd, 2020 at https://www.biorxiv.org/content/10.1101/2020.08.23.263327v1.

